# GexMolGen: Cross-modal Generation of Hit-like Molecules via Large Language Model Encoding of Gene Expression Signatures

**DOI:** 10.1101/2023.11.11.566725

**Authors:** Jiabei Cheng, Xiaoyong Pan, Yi Fang, Kaiyuan Yang, Yiming Xue, Qingran Yan, Ye Yuan

## Abstract

Designing de novo molecules with specific biological activity is an essential task since it holds the potential to bypass the exploration of target genes, which is an initial step in the modern drug discovery paradigm. However, traditional methods mainly screen molecules by comparing the desired molecular effects within the documented experimental results. The data set limits this process, and it is hard to conduct direct cross-modal comparisons. Therefore, we propose a solution based on cross-modal generation called GexMolGen (Gene Expression-based Molecule Generator), which generates hit-like molecules using gene expression signatures alone. These signatures are calculated by inputting control and desired gene expression states. Our model GexMolGen adopts a “first-align-then-generate” strategy, aligning the gene expression signatures and molecules within a mapping space, ensuring a smooth cross-modal transition. The transformed molecular embeddings are then decoded into molecular graphs. In addition, we employ an advanced single-cell large language model for input flexibility and pre-train a scaffold-based molecular model to ensure that all generated molecules are 100% valid. Empirical results show that our model can produce molecules highly similar to known references, whether feeding in- or out-of-domain transcriptome data. Furthermore, it can also serve as a reliable tool for cross-modal screening.

## Introduction

Designing novel molecules based on genetic information is a challenging but promising task [1]. Traditional search methods like CMap [2] compare observed gene expression changes, referred to as gene expression signatures, with those recorded in molecular perturbation experiments, which leads to a highly restricted search space that cannot accommodate the vast possibilities within the molecular space. Furthermore, they cannot effectively narrow down this search space by directly comparing cross-modal similarities.

Several existing *in silico* methods are attempting to overcome these shortcomings. Some models, like DLEPS [3], use an indirect approach by first predicting gene expression signatures as the drug effects and then using these signatures to screen molecules, as traditional methods do. Other methods, like WGAN [4] and Gex2SGen [5], directly input gene expression signatures and generate candidate molecules. WGAN uses generative adversarial networks (GAN), while Gex2SGen uses variational autoencoders (VAE). However, these models still need to improve their generalization ability. For example, they require input gene expression signatures that match the format used during the training phase; otherwise, retraining is required. Additionally, they mostly use a simplified molecular-input line-entry system (SMILES) to represent molecules. This form is a popular 1-dimensional representation that can be interpreted as specific “chemical text” so that it is easy to utilize advanced sequence networks like Transformer [6] fully. However, this representation form fails to preserve structural information and has limited potential to expand to higher dimensions [7]. On the other hand, the hypothesis behind this task is that molecules with induced similar gene expression signatures are likely to possess similar structures like scaffolds or pharmacophoric attributes. This structural information conflict makes it difficult to explore inherent mechanisms further [8]. Therefore, exploring new forms of modal encoders and modal interaction is necessary.

For molecular encoders, there are various graph-based approaches. For example, molecular autoencoders represent a molecule as a graph, where vertices correspond to atoms and edges represent chemical bonds [9, 10]. These methods intuitively capture the shape of molecules but need more efficiency due to low validity in generations. Another line of research involves flow-based methods like MolFlow [11], which achieve a perfect reconstruction rate but can be very computationally expensive for long chains. Scaffold-based approaches, such as hierVAE [12], utilize scaffolds as building blocks instead of individual atoms and edges, ensuring high validity and modest computational cost.

For genetic encoders, considering the remarkable success of foundation models that have revolutionized various fields, including omics data in life science, single-cell large language models hold great potential to be general encoders in this specific field. These models are pre-trained on large, unlabeled single-cell expression datasets and then fine-tuned for specific tasks [13]. High-quality models such as scBERT [14], Geneformer [15], TOSICA [16], scGPT [17], scFoundation [18], SCimilarity [19], and GeneCompass [20] are capable of receiving the flexible input form of gene expression signatures due to their powerful generalization abilities.

As to modal interaction, the “first-align-then-generate” strategy, inspired by models like DALL.E [21, 22], has gained great interest in biology. Recent work, including text-molecule matching [23], text-protein sequence generation [24], and PLIP [25], has achieved impressive performance.

Therefore, we propose GexMolGen, a cross-modal generation-based approach for designing molecules with specific biological activities.

## Materials & Methods

### Overview of GexMolGen

GexMolGen is a general framework capable of generating hit-like molecules from two specific gene expression inputs: initial and desired or post-perturbed states. The underlying principle is that molecular structure correlates with its resulting differential gene expression, also called gene expression signatures. GexMolGen adopts a strategy of “first-align-then-generate” for cross-modal generation. As shown in Fig.1A, the pipeline generally consists of 4 steps.

**Fig. 1.**
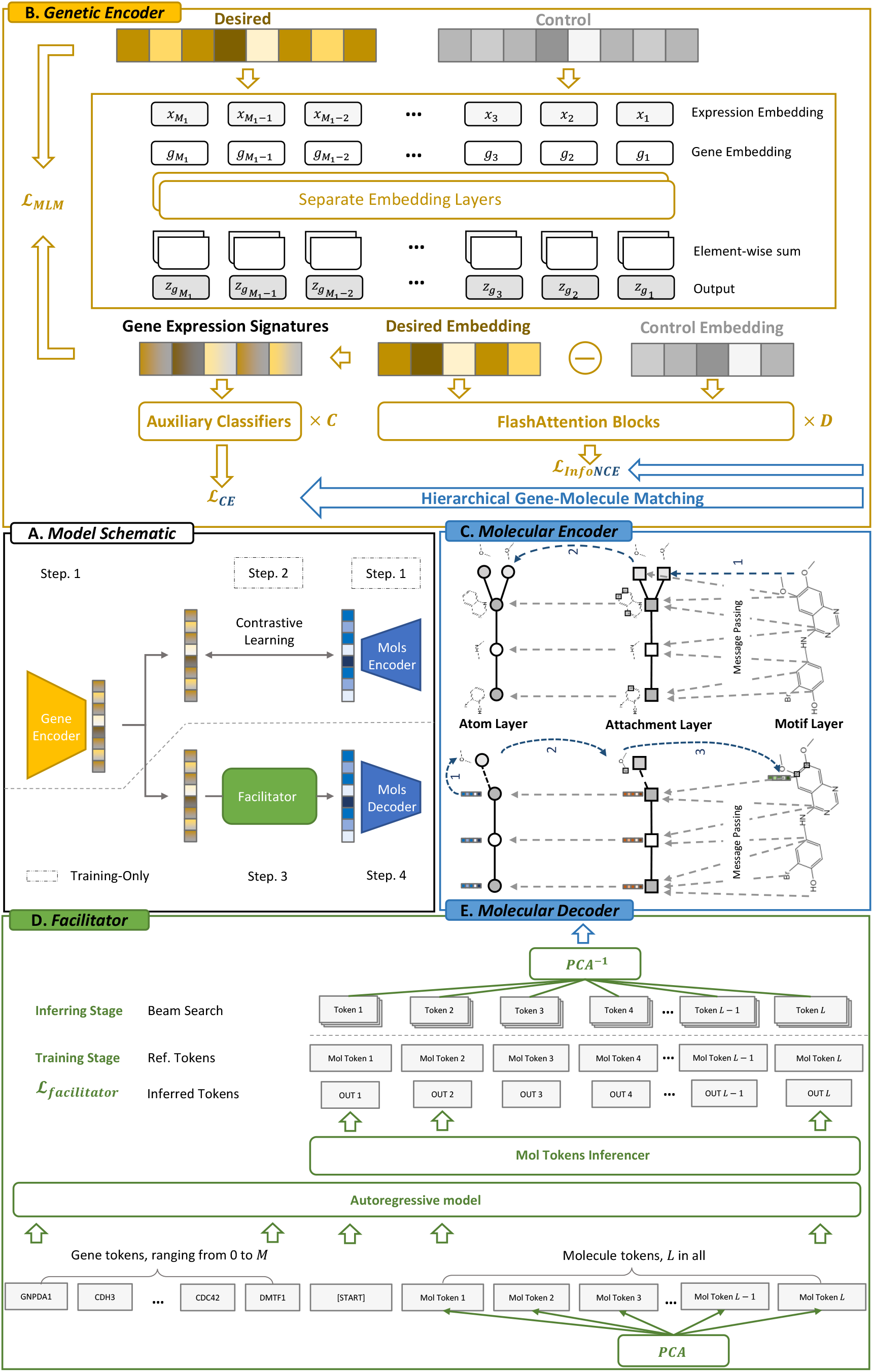
The workflow of GexMolGen employs a “first-align-then-generate” strategy (Section A), which consists of four steps: Step 1. Encoding of gene expression (Section B) and small molecule (Section C) data, respectively. Step 2. Aligning genetic modalities with molecular modalities (Section A). Step 3. Transformation from genetic modality to molecular modality (Section D). Step 4. Decoding to yield molecule graphs (Section E).

The 1st step involves the encoding of each modality. For the genetic encoder (visualized in Fig.1B), we apply an advanced single-cell foundation model, scGPT [17], to encode pre and post-perturbation gene expressions. We then calculate the difference within the gene embedding space to represent expression signatures. For the molecular encoder (illustrated in Fig.1C), we pre-train a scaffold-based molecular generator [12] on the ChEMBL [26] dataset and then use its encoder to produce molecular embeddings, thereby capturing structural information more efficiently than SMILES-based encoders [3, 4].

The 2nd step, as presented in Fig.1B, is to fine-tune each encoder through aligning. As complex negative examples are crucial to final alignment effects, we employ a hierarchical gene-molecule matching approach to enhance contrastive learning with iteratively refined granularity. The alignment projects gene expression signatures and molecules into a unified space, preparing for the transition from genetic embeddings to molecular ones.

The 3rd step, as depicted in Fig.1D, is implemented by a module entitled “Facilitator”, which is trained using a teacher-forcing technique and applied in an auto-regressive manner. Molecular embeddings are discretized into molecular tokens through Principal Component Analysis (PCA) and binning. Beam search is adopted in this module to optimize the diversity of the generation.

Finally, the inferred tokens are reordered and made continuous through retrieving and inverse PCA to molecular embeddings. They are then fed into the molecular decoder (as shown in Fig.1E) to generate molecular graphs. During the training phase, this module needs to compute the reconstruction loss to form a closed training loop within the framework.

The necessity for the molecular encoder and alignment phase is limited to the training phase only. For generation, the initial and the desired gene expressions and the number of molecular candidates are needed. For the retrieval function, additional molecules are required as a screening list. To our knowledge, GexMolGen is the first work that applies large language models to generate molecules cross-modally.

### Dataset Preprocessing

#### Dataset Used for Training

The datasets used in GexMolGen are derived from the L1000 public dataset (GEO ID: GSE70138) [27]. The L1000 technology uses microarray measurements to record the gene expression signatures caused by over 25,000 perturbations, including around 19,800 small molecules, in cell lines. Each perturbation is performed on multiple cell lines. This dataset is processed hierarchically and divided into five levels: Levels 1-3 record the absolute expression values of each gene in each bead, while Levels 4-5 report the differential expression of each gene obtained by comparing samples within each bead.

Specifically, we first collect the perturbed and control data from Level 3, ensuring they originate from the same bead. This data belongs to 6 major cell lines – VCAP, PC3, A549, A375, HT29, and MCF7. We average each molecule’s perturbed gene expressions, ignoring the cell line information, resulting in a final dataset size of 2×*N* ×*M*. Here, 2 signifies the experimental and control groups, *N* represents the number of molecules, and *M* indicates the count of gene symbols. Subsequently, we classify the structures based on the main layer of the InChI key, resulting in a total of 6360 classes. We randomly select 300 structures from this set, which include *N*_2_ types of small molecules, to form the hold-out set. Additionally, we choose another 300 structures, including *N*_3_ types of small molecules, to create the validation set. The remaining data is used for training the model with *N*_1_ types of small molecules.

We randomly sample *N*_*p*_ molecules from ChEMBL [26] as datasets for the molecular pre-training stage.

#### Dataset Used for Evaluating

In addition to the hold-out compound-induced gene expression data, we follow the same procedure as WGAN [4], our benchmarking method, to gather known inhibitors and genetic inputs for the comparative analysis of molecules generated from knock-out gene expressions. Initially, we select ten known inhibitors of human genes from the ExCape dataset [28]. These genes are AKT1, AKT2, AURKB, CTSK, EGFR, HDAC1, MTOR, PIK3CA, SMAD3, and TP53. Our selection criteria consist of a Pic50 value greater than or equal to 5, non-presence of the same InChi-key in the training set, and compliance with valence bond rules. Eventually, we collected 2,153, 1,293, 2,122, 1,609, 4,244, 1,939, 2,858, 2,292, 23,758, and 12,860 known inhibitors, the numbers respectively corresponding to each gene.

For each of these ten genes, we extract 100 transcriptome profiles from both the perturbed and control groups in the CRISPR knock-out perturbation data of the L1000 dataset [27].

### Molecular Encoder

For the molecular encoder (see Fig.1C), hierVAE [12] regards motifs as the smallest building blocks for generating small molecules. This approach achieves 100% validity in molecular generation. It encodes the input molecule hierarchically to form an atom layer 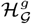, an attachment layer 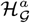, and a motif layer 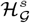. Each successive layer utilizes the previous one as its encoding condition.

#### Atom Layer

The input for the atom layer 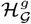 is the molecular graph *G*, the edge features {e (b_uv_)}, and the node features {e (a_u_)}. Through the message passing network [29, 30] *MPN*_*ψ*_(·), we obtain the encoded features:

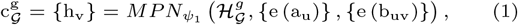

where h_v_ represents the updated atomic features.

#### Attachment Layer

The attachment layer 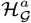 updates its nodes using the following equation:

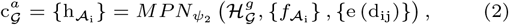

where e (d_ij_) denotes the edge features for the attachment layer, and 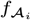 can be calculated based on atom vectors h_v_ and nodes *A*_*i*_ in the attachment layer:

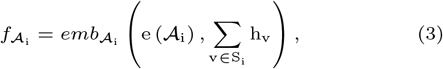

where 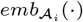 represents a fully connected layer and *e*(·) symbolizes a conventional embedding layer. The node *A*_*i*_ is the set of overlapping atoms between adjacent motifs.

#### Motif Layer

The motif layer 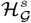 updates its nodes using the following equation:

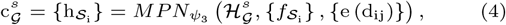

where e (d_ij_) denotes the edge features for the motif layer, and 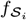 can be calculated based on 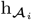 and *S*_*i*_:

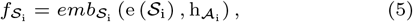

where 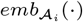 represents a fully connected layer and *e*(·) symbolizes a conventional embedding layer. Nodes *S*_*i*_ in the motif layer represent the type of the motifs.

Finally, the embedding of the *i*-th molecule is defined as:

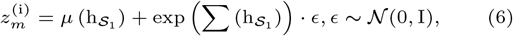

where *µ*(·), exp(·) are both fully connected layers representing the mean and log variance, respectively, and *S*_1_ represents the root motif, which is also the first motif for decoding.

### Molecular Decoder

During decoding (see Fig.1E), we continue to adopt the molecular embedding notion as *z*_*m*_, and the motif layer encoded vectors as 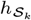, where *k* signifies the step. Initially, we set *k* = 1. Then, the hierarchical decoding of the molecular graph is shown as follows.

#### Motif Prediction

This step predicts the next motif *S*_*t*_ to be attached to *S*_*k*_. It is implemented as a classification task over the motif vocabulary *V*_*S*_ :

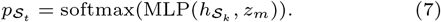

#### Attachment Prediction

Similar to motif prediction, but applying to the attachment vocabulary *V*_*A*_(*S*_*t*_), this step predicts the attachment configuration between *S*_*t*_ and *S*_*k*_. In other words, it determines the type of atom that connects *S*_*t*_ and *S*_*k*_:

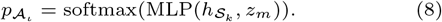

#### Graph Prediction

This step is to decide how *S*_*t*_ will be attached to *S*_*k*_. After attachment prediction, we only narrow down to the connected atom type but do not specify the exact position. The attachment between *S*_*t*_ and *S*_*k*_ is defined as atom pairs ℳ_*tk*_ = {(*u*_*j*_ .*v*_*j*_)|*u*_*j*_ ∈ *A*_*k*_, *v*_*j*_ ∈ *A*_*t*_} where atom *u*_*j*_ and *v*_*j*_ are attached together. The probability of a candidate attachment, *M*, is calculated based on the atom vectors, 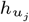 and 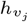 :

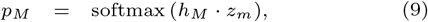

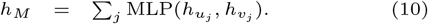

#### Training

During training, this module needs to minimize the negative ELBO:

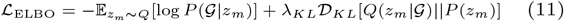

to complete the training loop.

#### Validation

During validation, we utilize the Fréchet ChemNet Distance (FCD) [31] as a selection criterion for models. This metric evaluates whether the generated molecules show diversity and possess chemical and biological properties similar to reference molecules.

### Genetic Encoder

In Fig.1B, a single-cell large language model known as scGPT [17] is employed as the genetic encoder. Trained on an extensive amount of unlabeled single-cell expression data, this model shows potential in capturing underlying biological signals. The pretrained gene tokens from scGPT can uniquely identify each gene in the input data, allowing for flexible gene inputs during the encoding process. Our model receives two sets of gene expression data: the initial and the desired or post-perturbation data. Both sets of data independently undergo the same encoding pipeline. Using the *i*-th gene expression data point as an example, we initially retrieve its corresponding gene tokens from the scGPT pretrained gene dictionary:

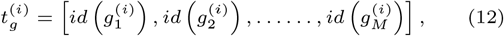

where 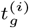 represents the gene token encoding for the *i*-th data, and 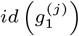 represents the unique integer identifier for gene *g*_*j*_ in the *i*-th data.

Next, we preprocess the expression values by implementing transcripts per thousand (TPT) normalization and *log*1*p* transformation:

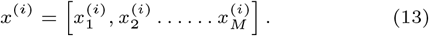

The gene embedding is then defined as follows:

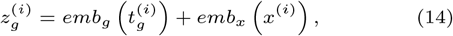

where *emb*_*g*_ (·) refers to the conventional embedding layers, and *emb*_*x*_ (·) represents the fully connected layer.

We further utilize the mask language model (MLM) task to fine-tune scGPT. Instead of commonly using mean square loss (MSE), we use the autofocus direction-aware loss in GEARS [32]. This loss automatically assigns a greater weight to differentially expressed genes by amplifying the exponent of the error. Given a minibatch of *B*_1_ data points, we randomly sample 95%

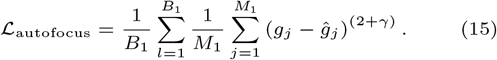

Moreover, we introduce an additional direction-aware loss to increase the predicted data’s sensitivity to the direction relative to the control group, which is defined as follows:

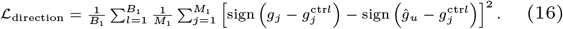

The optimization objective for the MLM task, denoted as ℒ_MLM_, is ultimately calculated by taking a weighted sum of the values of ℒ_autofocus_ and ℒ_direction_:

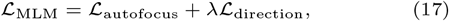

where *λ* functions to adjust the weight for the directional loss.

We then represent the gene expression signatures for subsequent alignment by calculating the difference in post-encoding embedding between the desired and control states:

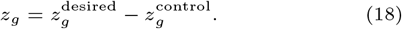

### Alignment

Alignment is the 2nd step (see Figure.1A) and aims to extract the mutual information between genetic and small molecular modalities, thus enhancing the controllability of the generation model. In this process, we adopt contrastive learning [33, 34] for multi-modal matching and introduce a hierarchical gene-molecule matching strategy to overcome model collapse.

#### Hierarchical clustering

Hierarchical clustering merges data into clusters step by step by calculating their similarity until a complete tree-like structure is formed. We use this strategy to cluster the small molecule embeddings into *C* = {*c*_1_, *c*_2_, *c*_3_} classes. The detailed process is shown in Algorithm 1.

#### Hierarchical Gene-Molecule Matching

Through hierarchical clustering, molecules are grouped into clusters, and the model can be guided by the cross-entropy loss function at a coarse resolution. Given that the small molecules have been clustered into *C* = *{c*_1_, *c*_2_, *c*_3_} classes, we require the genetic encoder to classify the corresponding perturbations based on gene expression signatures. The cross-entropy loss function measures the quality of this task:

##### Algorithm 1 Hierarchical Clustering

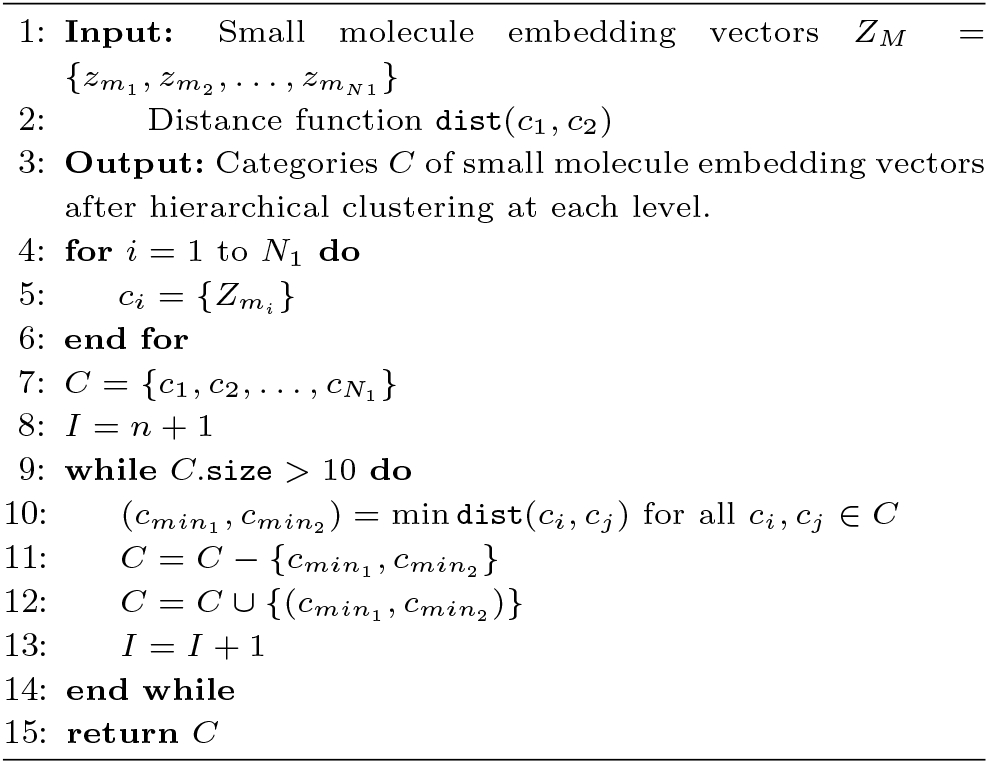

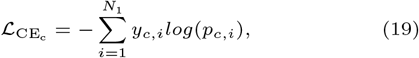

where *y*_*c,i*_ and *p*_*c,i*_ denote the truth labels and predicted labels for the *c*-th cluster of small molecules based on gene embeddings.

#### Contrastive Learning Alignment

The final layer *z*_*g*_ output in the genetic encoder is aligned with small molecular vectors *z*_*m*_ via contrastive learning. The optimization objective is to minimize the InfoNCE loss function [35]:

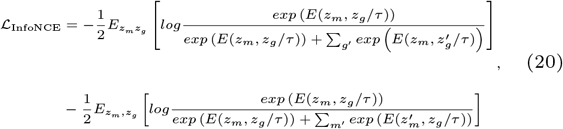

where *z*_*m*_^*′*^ and *z*_*g*_^*′*^ are unpaired molecule-gene negative sample pairs in a minibatch composed of *B*_2_ data points, *E*(.,.) is an energy (scalar) function defined on these two modalities. We use vector dot product for representation; *τ* is a learnable coefficient. The InfoNCE loss aims to minimize the distance between positive samples and maximize the distance between negative samples.

Overall, during the learning process, our genetic encoder optimizes two functions: InfoNCE loss ℒ_InfoNCE_ for contrastive learning, and cross-entropy loss 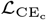 for the hierarchical gene-molecule matching, which can also serve as the hard sample mining task:

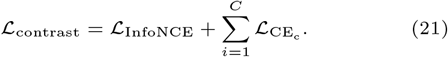

### Facilitator

Assuming our dataset consists of sample pairs (*m, g*) where *m* represents a small molecule and *g* represents its corresponding gene expression signatures, we model the transition from the gene expression signatures to the molecules, denoted as *P* (*m*|*g*), using a two-phase process involving modality facilitation and model generation [36]. Let *z*_*m*_ and *z*_*g*_ continue representing the small molecule and gene embeddings, respectively. Thus, *P* (*m*|*g*) can be decomposed as

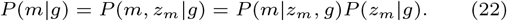

The first equation holds because *z*_*m*_ is a deterministic function of the small molecule dataset *m*, and the second equation holds due to the chain rule [22, 21].

When the small molecule decoder *P* (*m*|*z*_*m*_, *g*) has been determined [12], the primary focus is on optimizing the facilitator *P* (*z*_*m*_|*g*) (see Fig.1D). We adopt an auto-regressive model for modality transformation [37, 38, 39].

Since *z*_*m*_ is a continuous value, the cross-entropy loss function cannot be directly applied. To solve this, we adopt the approach utilized in DALL·E 2 [22], where we reduce the dimension of *z*_*m*_ from *d* to *L* using principal component analysis (PCA). This reduction allows us to retain nearly all the information (with a reconstruction mean squared error below 1%). Subsequently, we discretize the *L*-dimensional values into *N*_*m*_ bins, assuming that these values are uniformly distributed.

Fig.1D provides the mechanism of the Facilitator. It takes as input *M* genes, a special symbol [start], and *L* small molecule tokens and sequentially produces inferred tokens 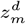. The loss function is defined as follows:

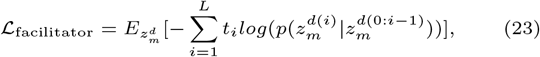

where *t*_*i*_ represents the *i*-th true token, and 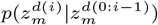 represents the inferred token of the small molecules.

#### Beam Search Optimization

Beam search is a commonly used search algorithm in sequence generation tasks [40, 9]. It utilizes dynamic programming principles by keeping track of a set of candidate sequences, known as the beam, with a beam width of *K* to preserve promising results. The beam search algorithm effectively manages the search space and improves the accuracy and diversity of generated sequences. Specifically, the beam search algorithm follows the following steps:

1. Initialize the beam with a set of candidate sequences, each consisting of *M* gene tokens and [start].
2. At each time step:
  a. Calculate the likelihood of every candidate token becoming the next token based on the current sequence.
  b. Select the top *K* candidate sequences based on their scores, where *K* is the beam width.
  c. Expand each selected candidate sequence by appending the molecular tokens and updating its score.
3. Repeat step two until a complete sequence of small molecule tokens *L* is formed.
4. Return the top-ranked sequence from the beam as the final output.

After the beam search process, the final output tokens 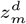 are transformed into *z*_*m*_ using the inverse of PCA. The continuous *z*_*m*_ values are then fed into the molecular decoder *P* (*m*|*z*_*m*_, *g*) [12]. This decoding procedure results in *K* sets of molecule candidates.

## Results

### Generating Hit-like Molecules from Compound-Induced Gene Expression Signatures

In this section, we assess the performance of our model GexMolGen in generating hit-like molecules using compound-induced gene expression signatures. We input a set of *N*_3_ hold-out samples into the model and compare the molecules generated with the reference ones in hold-out samples.

In Fig.2, we illustrate this comparison (for more details, please refer to Table A.1). The results indicate that the generated molecules can successfully reproduce some complex scaffold structures that appeared in the reference molecules, such as long chains (see Fig.2(3)(6)), cyclic structures (see Fig.2(1)), and cross-like structures (see Fig.2(7)(9)).

**Fig. 2.**
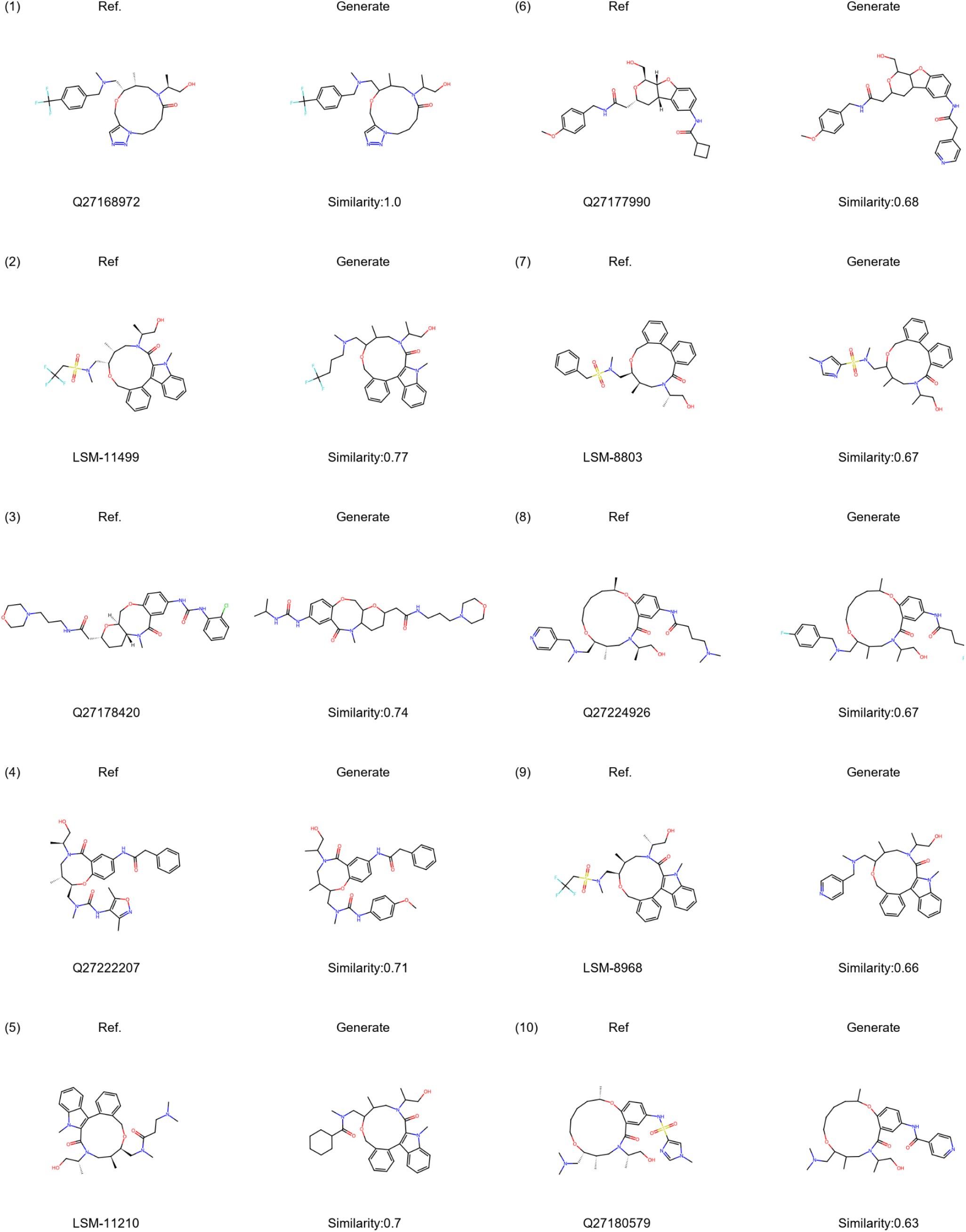
Examples of generated molecules using a compound-induced gene expression signature. Compared to reference small molecules, this figure showcases examples of hit-like small molecules generated by GexMolGen from compound-induced gene expression signatures. The similarity calculation metric adopts Tanimoto’s similarity using Morgan fingerprints.

We compare our model with two traditional methods: searching molecules in the training set according to the cosine similarity or Euclidean distance among gene expression signatures [4]. We also include a random model that generates directly from samples in the molecular latent space to demonstrate the guided influence of genetic information. We adopt Tanimoto similarity based on Morgan fingerprints (radius=3, 1024 bits) as the global similarity metric and use Tanimoto similarity based on MACCS keys and Fraggle similarity as the local structural metric.

Fig.3 shows the distribution of these metrics for different methods. Detailed results can be found in Table A.1, A.2, A.3. The results obtained from our model GexMolGen demonstrate a significant improvement over the other three baseline methods (p-value <0.001, according to a one-sided Mann-Whitney U-test). Specifically, our model achieves a mean Fraggle score of 0.57, a mean Morgan-based Tanimoto score of 0.30, and a mean MACCS-based Tanimoto score of 0.70. These results indicate that the molecules generated by our model exhibit high similarity to the reference molecules in both fingerprints and substructures. Moreover, across all three metrics, the Euclidean distance search method is consistently inferior to the cosine similarity method. This discrepancy can be attributed to the curse of dimensionality, where the Euclidean distance becomes less sensitive to differences in gene expression signatures as the dimension increases. We also notice that the randomly generated model has the poorest overall performance. Although it occasionally produces molecules with high similarity scores (e.g., 1.19% with a similarity >0.29 in Fig.3(b)), it also generates a more significant proportion of irrelevant molecules (e.g., 26.27% with a similarity <0.08 in Fig.3(b)), which highlights the significance of guidance from gene expression signatures in generating desired molecular properties.

**Fig. 3.**
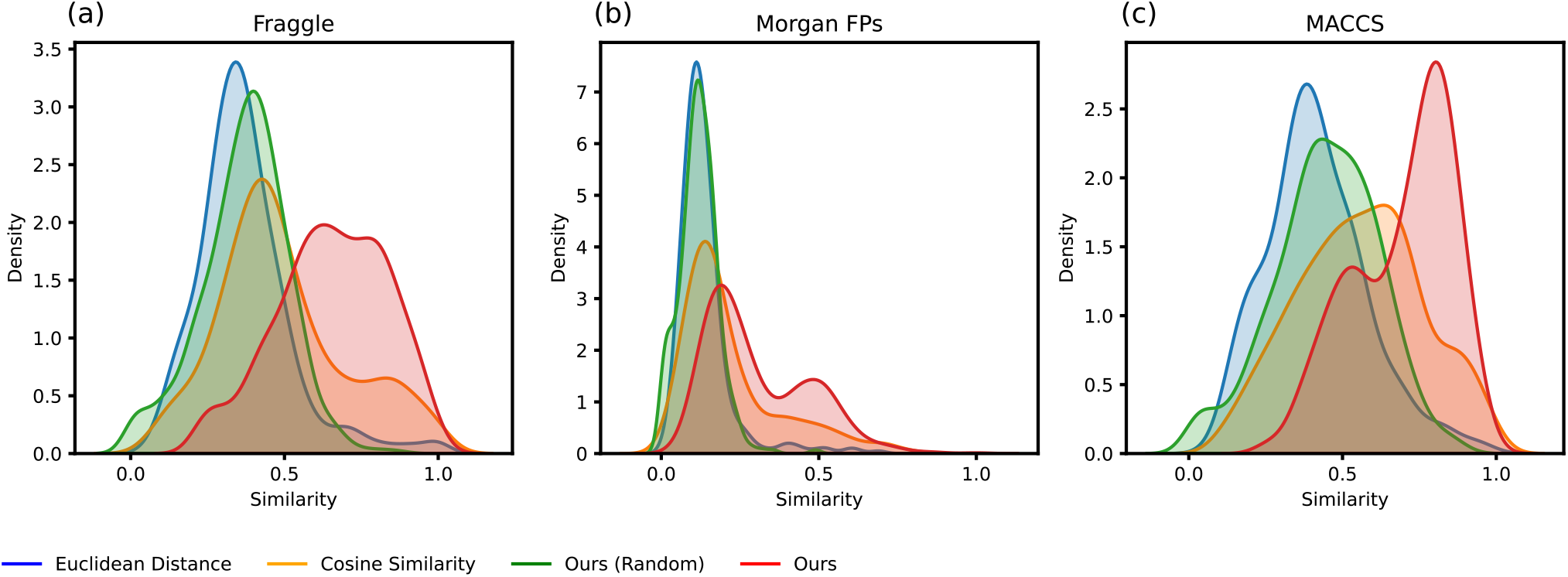
Distribution of similarity. This figure shows the comparison results in generating hit-like molecules from compound-induced gene expression signatures. (a), (b) and (c) represent the distributions of metrics calculated using Fraggle similarity, Morgan fingerprints, and MACCS keys, respectively. The comparison methods include traditional searching methods such as cosine similarity and Euclidean distance, as well as the random model and ours, each represented by different colors. Supporting materials can be found in Table A.1, A.2 and A.3.

### Generating Hit-like Molecules from Knock-out Gene Expression Signatures

To test the generalization ability of our model GexMolGen, we evaluate it to generate potential gene inhibitors from knock-out gene expression data, which can also serve as an out-of-distribution test. The underlying hypothesis is that inhibitors of a specific gene will cause equivalent changes in gene expression when that gene is knocked out.

We utilize the same evaluation metrics and comparison methods described in the Result 3.1. The collection of knock-out data and the corresponding known inhibitors can be found in the Methods 2.2.2. The procedure involves each method generating 100 candidate molecules for each gene based on the given knock-out transcriptome. Subsequently, we record the scores between each candidate and its most similar known inhibitors.

Fig.4 and Fig.5 presents a visualization of the molecules generated by our model and the most similar known inhibitors. Morgan-based Tanimoto similarity scores measure this similarity. Compared to WGAN [4] with an average score of 0.26 (see Table A10 for details), our model generates molecules that not only exhibit a higher overall degree of similarity to known inhibitors but also successfully reconstructs the continuous complete skeletal structure of the molecules.

**Fig. 4.**
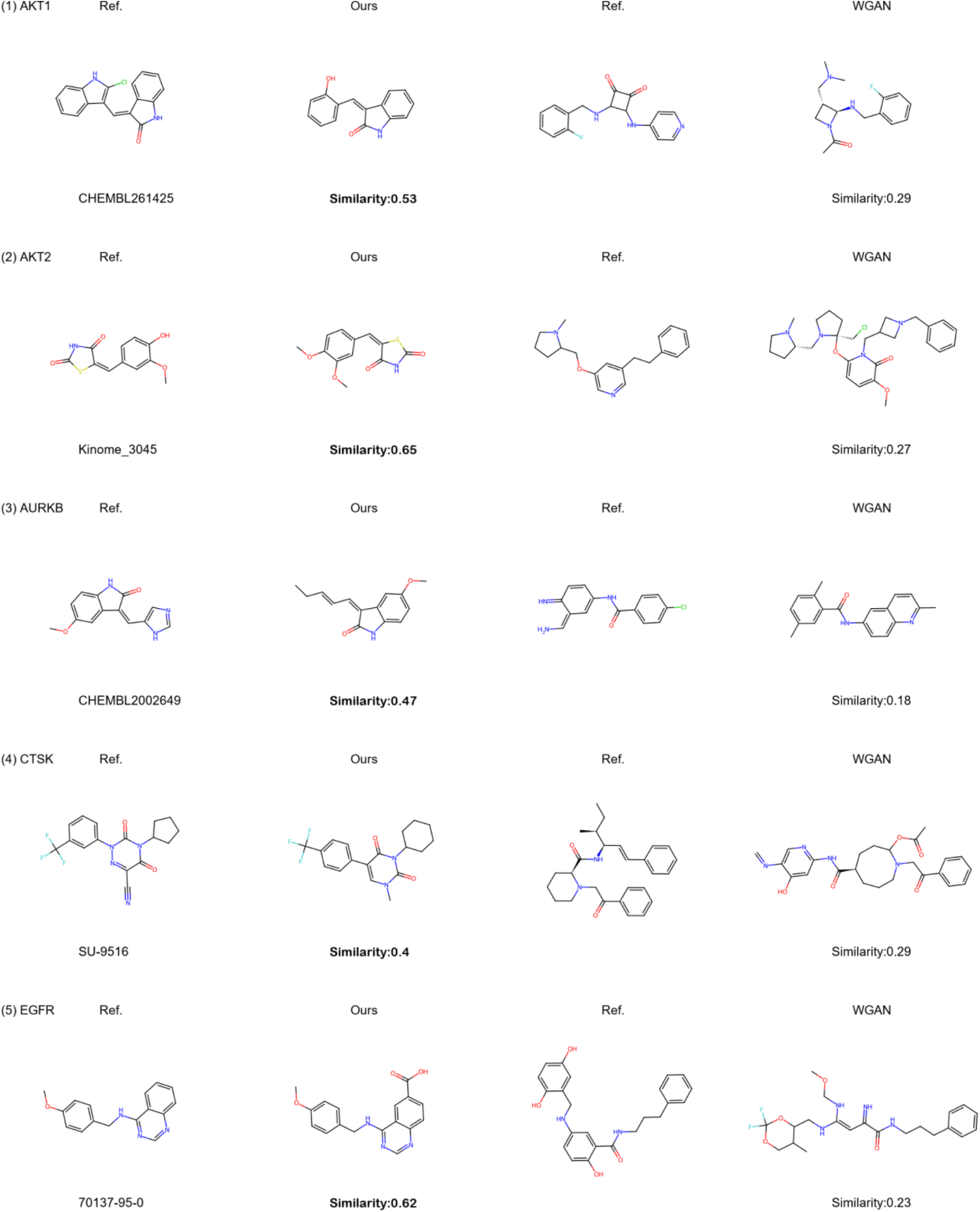
Examples of generated molecules using knock-out gene expression signatures. This figure compares our method and the results generated by WGAN [4]. The title is the name of the knock-out gene, the 1st and 3rd columns are the most similar known inhibitors each found, and the 2nd and 4th columns are the outcomes generated by the two methods. The results of WGAN are extracted from Figure 2 of the paper using OCR, and the specific results can be referred to in Table A10.

**Fig. 5.**
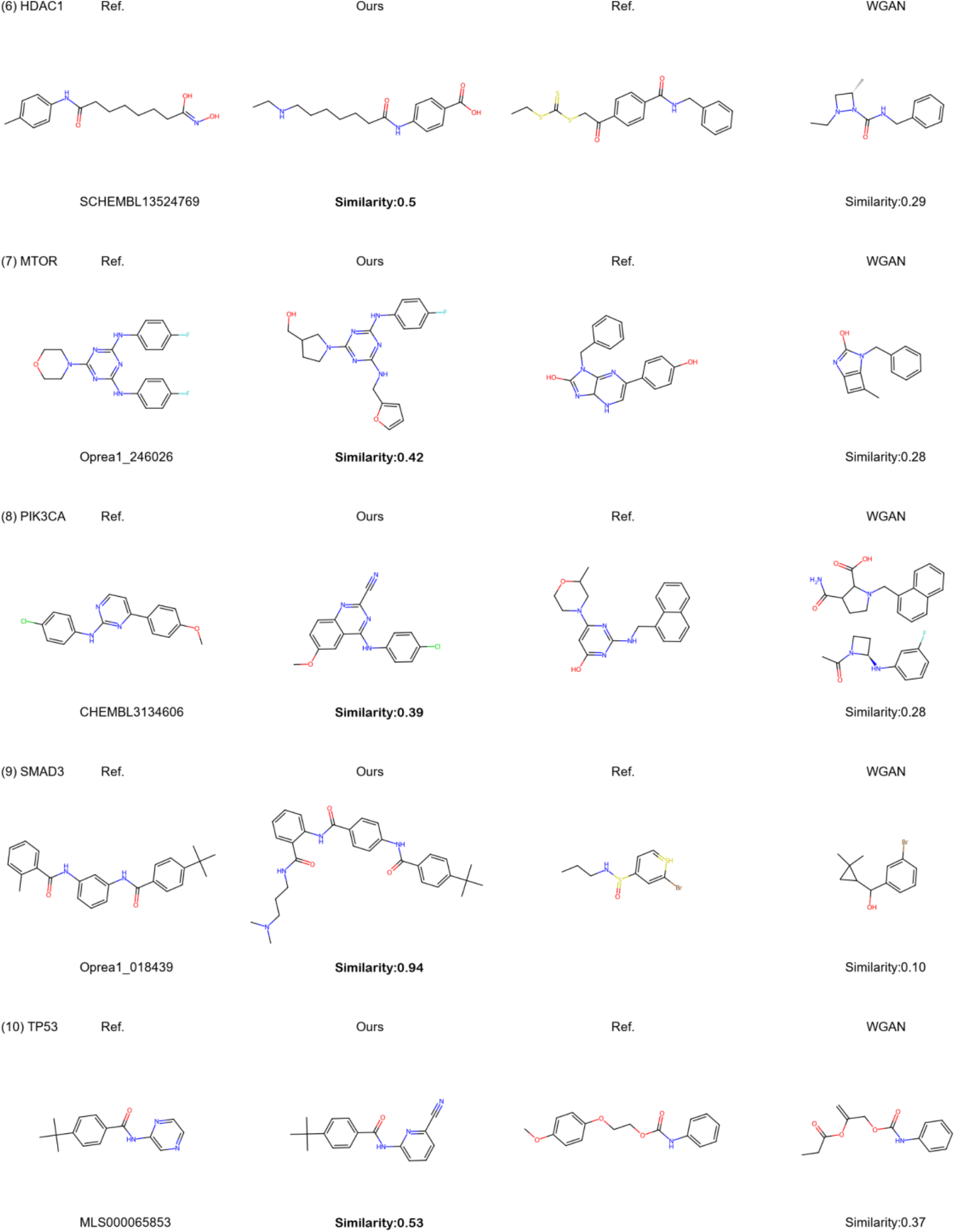
Examples of generated molecules using knock-out gene expression signatures. This image is an extension of the results from Fig.4.

The results of the experiments are thoroughly demonstrated in Fig.6. Generally, our model significantly surpasses the three other baseline methods with higher average scores and diversity in most cases. Furthermore, due to the limited size of the search database, the molecule candidates of traditional methods exhibit shallow diversity. For instance, CTSK, in MACCS-based metrics, shows a nearly linear form. On the other hand, the random method produces the widest range of scores due to the unconditional molecular sampling, which simultaneously leads to the deterioration of molecular properties. It occasionally produces molecules with high similarity scores but generates many molecules with very low correlations to known inhibitors, as observed in Fig.6’s AKT2. By leveraging genetic information, our model narrows down the generation space and allows for some diversity, striking a balance between structure similarity and molecular diversity. We also refer to how other deep learning methods perform on similar tasks, as presented in Table 1. Despite some differences in the tested genes, it can be observed that our model consistently produces competitive results. The notable advantages of our model are its efficiency with 100% generated molecular validity and high input flexibility, which are missing in other methods. The details of the results can be found in Table A.4, A.5, A.6.

**Fig. 6.**
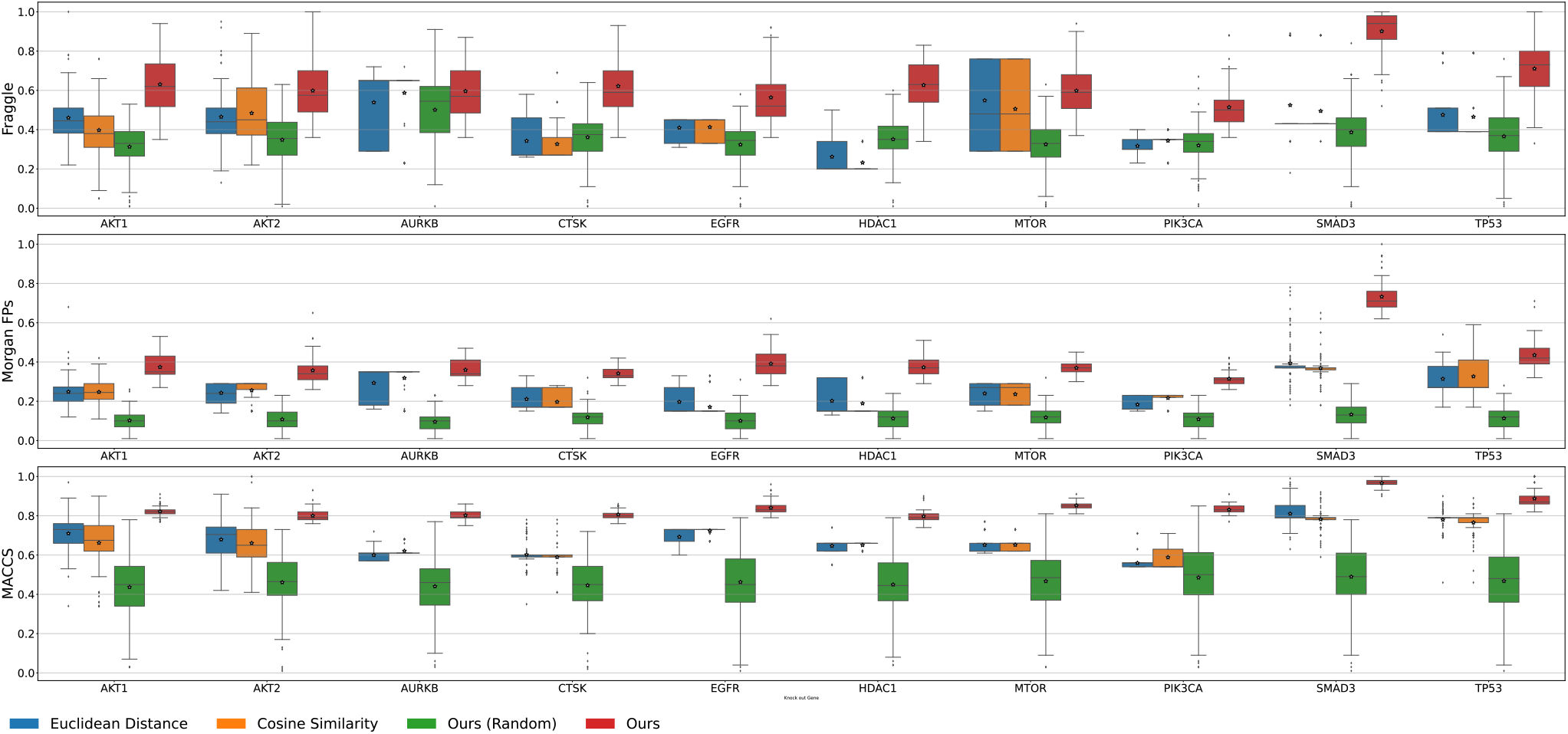
Distribution of similarity. This figure demonstrates the comparison results of potential gene inhibitors generated from knock-out gene expression signatures. The y-axis denotes three metrics: Fraggle similarity, Morgan fingerprints, and MACCS keys. The x-axis denotes the 10 test genes. Different colors represent different methods. The hollow pentagrams represent the mean values. Supporting materials can be found in Table A.4, A.5, and A.6.

**Table 1.** Metrics comparison with other deep learning methods. Comparing GexMolGen with other deep-learning approaches in generating potential inhibitors from knock-out transcriptomes.

| Name<sup>1</sup> | Data<sup>2</sup> | Form<sup>3</sup> | Valid<sup>4</sup> | Metric<sup>5</sup> |
| --- | --- | --- | --- | --- |
| WGAN [4] | L1000 | SMILES | 10% | 0.15 ~ 0.70 |
| Bidirectional Adversarial AE [41] | L1000 | SMILES | 76% | 0.30 ~ 0.48 |
| TRIOMPHE [42] | L1000 | SMILES / SELFIES | 16.89%/ 93.96% | 0.29 ~ 0.68 |
| PaccMann [43] | GDSC/CCLE | SMILES | 96.2% | 0.27 ~ 0.45 |
| Deep Generative with RL [44] | TCGA project | SMILES | 95.9% | 0.47 ~ 0.83 |
| Ours | L1000 | Graph | **100%** | **0.72 ~ 1.00** |

<sup>1</sup>The 1st column (Name) outlines the name and source of these methods.
<sup>2</sup>The 2nd column (Data) is the types of datasets.
<sup>3</sup>The 3rd column (Form) specifies the representation of molecules.
<sup>4</sup>The 4th column (Valid) reports the average validation of their molecular generator.
<sup>5</sup>The 5th column (Metric) outlines the range of Tanimoto similarity.

### Screening Molecules Using Cross-Modal Similarity

Our model GexMolGen also offers a well-defined cross-modal similarity for measuring the correlation between gene expression signatures and molecular structures. Therefore, it also supports screening molecules using gene expression profiles alone.

We evaluated the effectiveness of GexMolGen by the task described in Result 3.2. Specifically, our model first encodes the molecules from the reference dataset into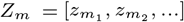, and the gene expression data into 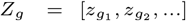 respectively, then calculates their alignment scores *Z*_*m*_^*T*^ · *Z*_*g*_ to find the molecule *m*_·_ with the highest score for each genetic input *g*_·_. The visualization of the screening pipeline can refer to Fig.7. This procedure is similar to the operations in traditional search methods, but the latter only calculates similarities within a uni-modal scope. For instance, in CMap [2], gene expression queries are compared with recorded gene expressions that molecules have perturbed to analyze modal similarity.

**Fig. 7.**
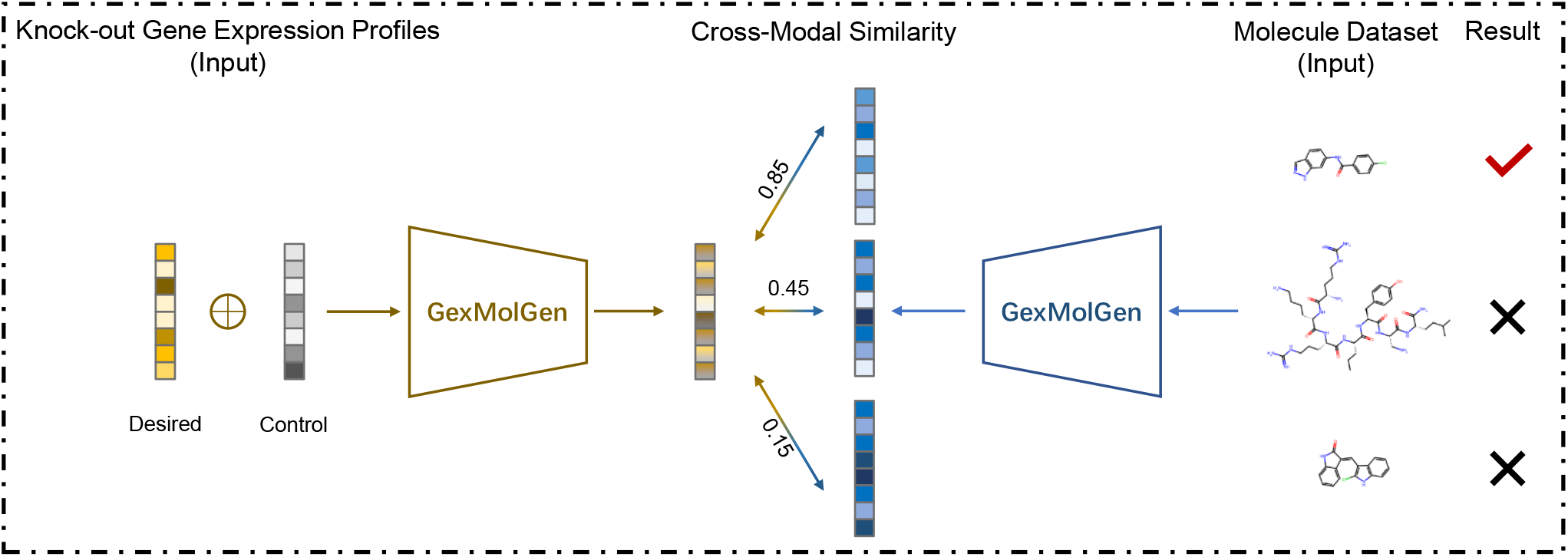
The flowchart using only our model’s retrieving function.

Fig.8 displays some examples of screening from the same molecule datasets, and Table 2 demonstrates statistics. Both indicate that the molecules screened using our cross-modal similarity are much more similar to known inhibitors than those found by traditional methods. This result reflects the advantage of this direct cross-modal similarity calculation. The supporting materials are given in Table A.7.

**Fig. 8.**
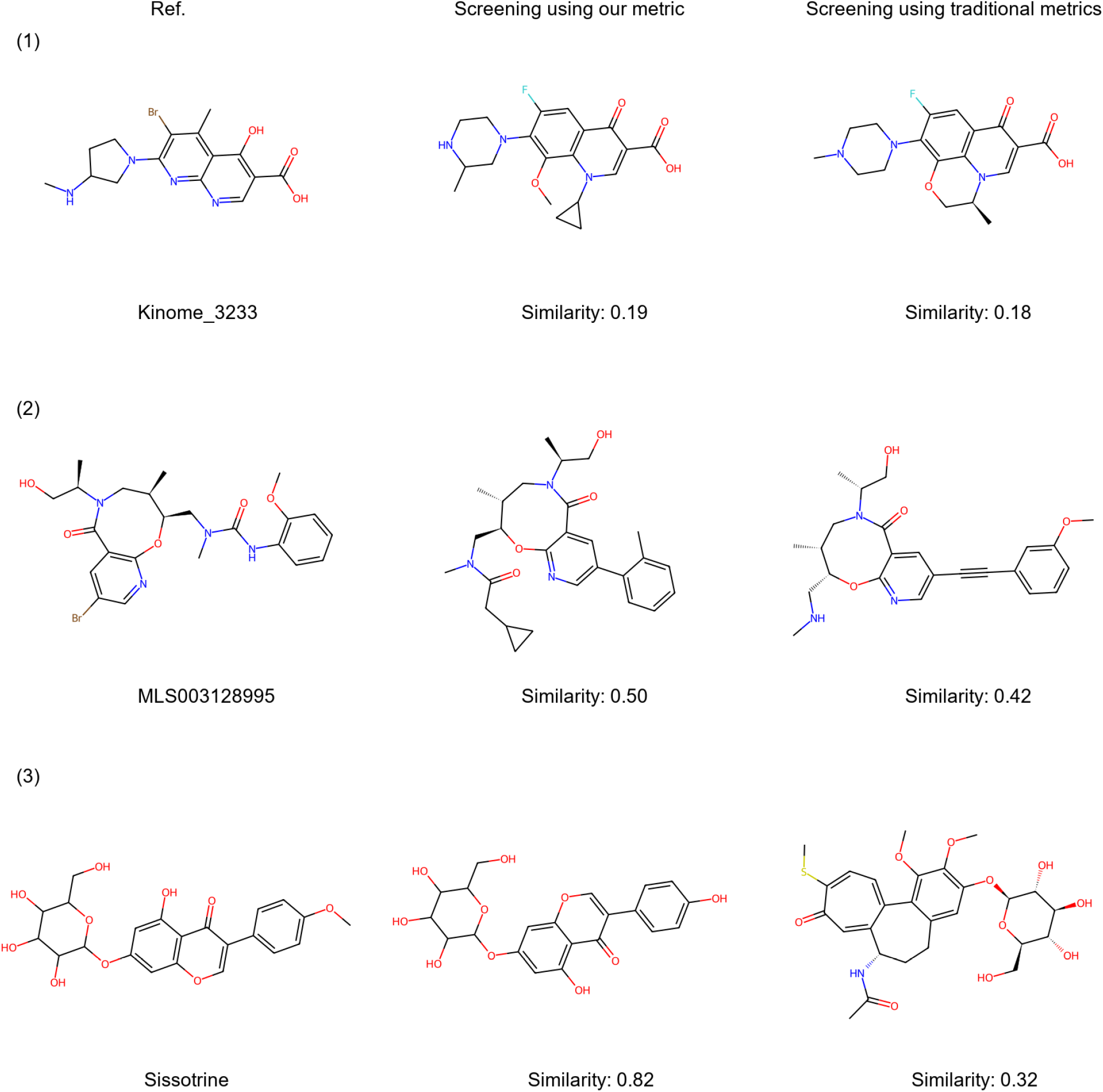
Examples of searched molecules using knock-out gene expression signatures. This figure presents instances of molecules separately screened by our metric and traditional metrics but share the most similar inhibitors. Detailed results are in Table A.7.

**Table 2.** Summary statistics of similarity. This table presents the results of searching potential inhibitor using our cross-modal similarity metric and the indirect gene similarity calculation metric using traditional methods. The results reported are Tanimoto similarities calculated by MACCS fingerprints, and the highest values are highlighted in **bold**. Results under the other two similarity metrics can be found in Table A8 and A9.

| Knock-out Genes | MACCS Scores (Ours) | MACCS Scores (Cos.) | MACCS Scores (Euc.) |
| --- | --- | --- | --- |
| AKT1 | **0.745 ± 0.056** | 0.710 ± 0.091 | 0.662 ± 0.130 |
| AKT2 | **0.690 ± 0.065** | 0.679 ± 0.098 | 0.661 ± 0.113 |
| AURKB | **0.694 ± 0.047** | 0.600 ± 0.024 | 0.621 ± 0.025 |
| CTSK | **0.658 ± 0.086** | 0.603 ± 0.069 | 0.590 ± 0.054 |
| EGFR | **0.752 ± 0.054** | 0.693 ± 0.051 | 0.724 ± 0.017 |
| HDAC1 | **0.649 ± 0.058** | 0.647 ± 0.028 | 0.651 ± 0.017 |
| MTOR | **0.740 ± 0.087** | 0.652 ± 0.045 | 0.652 ± 0.035 |
| PIK3CA | **0.684 ± 0.053** | 0.560 ± 0.030 | 0.588 ± 0.070 |
| SMAD3 | **0.820 ± 0.069** | 0.811 ± 0.065 | 0.782 ± 0.050 |
| TP53 | **0.793 ± 0.061** | 0.780 ± 0.055 | 0.766 ± 0.062 |

## Discussion

This paper proposes GexMolGen, a method to generate hit-like molecules based on signatures calculated by desired gene expression relative to the initial states. We adopt a “first-align-then-generate” [21] strategy for cross-modal generation. Specifically, we utilize an advanced single-cell large language model called scGPT [17] as the genetic encoder and pre-train hierVAE [17], a graph-based molecular model. To enhance the effects of contrastive learning, we integrate a hierarchical gene-molecule matching approach for alignment, serving as the mining task of hard negative samples. The facilitator initially transitions genetic embeddings into molecular tokens during the generation process. These tokens are subsequently transformed into a continuous molecular embedding, fed into the decoder to generate hit-like molecules.

Our method demonstrates competitive and superior performance compared to traditional search methods in generating molecules from in-domain compound-induced expression signatures and out-of-domain gene knock-outs. Additionally, our model exhibits efficiency by ensuring 100% validity and supports input flexibility, surpassing other deep-learning methods. It also can be used as a screening tool since it provides an explicit metric for measuring the similarity between genetic and molecular modalities.

## Key Points

- We propose a novel cross-modal generation-based approach for designing molecules with specific biological activities.
- GexMolGen is general and accepts flexible input formats by fully leveraging the power of foundation models.
- GexMolGen has an explicit metric to measure the correlation between inputs and outputs and retrieval.
- GexMolGen has 100% validity because of its scaffold-based molecular generation strategy.

## Declarations

### Code Availability

The source codes of GexMolGen are available on Zenodo (https://zenodo.org/records/11092780?token=eyJhbGciOiJIUzUxMiIsImlhdCI6MTcxNDQ4MjYwOCwiZXhwIjoxNzk4Njc1MTk5fQ.eyJpZCI6IjQwMjc0ZTlkLTQ0YjItNDY0Mi1hYmRjLTlkM2VkMjVhNWE5OSIsImRhdGEiOnt9LCJyYW5kb20iOiI5MzhjMjdmZjdmNmRlZmMxZTU0MTNhMGNmZGIzMDNmOCJ9.p3lxB8h_BDhuHm_1yqr4L1ZjRCTXmftMzpNdYaexmvycISDkoUu1cizUFrmcRZ52QovkPeQDjsHBrdScRkVqiw.), together with a usage documentation and setup environment.

### Author contributions

J.C., Q.Y., and Y.Y conceived the concept of this work. The experiments were performed by J.C., with the assistance of X.P., Y.F., K.Y., and Y.X. All authors participated in analyzing the results and writing the manuscript. The project was under the supervision of X.P., Q.Y. and Y.Y.

### Data Availability Statements

The data underlying this article are available in the article and its online supplementary material. The curated data can be found at https://zenodo.org/records/11100665?token=eyJhbGciOiJIUzUxMiIsImlhdCI6MTcxNDYyNzUzOSwiZXhwIjoxNzk4Njc1MTk5fQ.eyJpZCI6IjM3ZmFkNGU4LWViMDYtNGNkNy1iOTc4LWI0ZTBkMDk2OWI0YyIsImRhdGEiOnt9LCJyYW5kb20iOiJhOGJjZDM5NWFkY2ZiNDAwNjAzYzIwMTg2ODNjYWI2NCJ9.dADyS-0PBsFKr_z1yDdcDnoGoY5PFOSbnYtt6aIz4RLoNxykoIQffAlzQDPbFgqnZJmp7PmjNXPCXHkDMuZHuA. The raw data is downloaded from https://clue.io/data/CMap2020#LINCS2020, https://www.ebi.ac.uk/chembl/ and https://solr.ideaconsult.net/search/excape/#.

## References

1. Fabien Vincent, Arsenio Nueda, Jonathan Lee, Monica Schenone, Marco Prunotto, and Mark Mercola. Phenotypic drug discovery: recent successes, lessons learned and new directions. Nat Rev Drug Discov, 21(12):899–914, December 2022. Number: 12 Publisher: Nature Publishing Group.

2. Aliyu Musa, Laleh Soltan Ghoraie, Shu-Dong Zhang, Galina Glazko, Olli Yli-Harja, Matthias Dehmer, Benjamin Haibe-Kains, and Frank Emmert-Streib. A review of connectivity map and computational approaches in pharmacogenomics. Briefings in Bioinformatics, 18(5):903, September 2017.

3. Jie Zhu, Jingxiang Wang, Xin Wang, Mingjing Gao, Bingbing Guo, Miaomiao Gao, Jiarui Liu, Yanqiu Yu, Liang Wang, Weikaixin Kong, Yongpan An, Zurui Liu, Xinpei Sun, Zhuo Huang, Hong Zhou, Ning Zhang, Ruimao Zheng, and Zhengwei Xie. Prediction of drug efficacy from transcriptional profiles with deep learning. Nat Biotechnol, 39(11):1444–1452, November 2021. Number: 11 Publisher: Nature Publishing Group.

4. Oscar Méndez-Lucio, Benoit Baillif, Djork-Arné Clevert, David Rouquié, and Joerg Wichard. De novo generation of hit-like molecules from gene expression signatures using artificial intelligence. Nat Commun, 11(1):1–10, January 2020. Number: 1 Publisher: Nature Publishing Group.

5. Dibyajyoti Das, Broto Chakrabarty, Rajgopal Srinivasan, and Arijit Roy. Gex2SGen: Designing Drug-like Molecules from Desired Gene Expression Signatures. J. Chem. Inf. Model., 63(7):1882–1893, April 2023. Publisher: American Chemical Society.

6. Ashish Vaswani, Noam Shazeer, Niki Parmar, Jakob Uszkoreit, Llion Jones, Aidan N. Gomez, Łukasz Kaiser, and Illia Polosukhin. Attention is all you need. In Proceedings of the 31st International Conference on Neural Information Processing Systems, NIPS’17, pages 6000–6010, Red Hook, NY, USA, December 2017. Curran Associates Inc.

7. Maranga Mokaya, Fergus Imrie, Willem P. van Hoorn, Aleksandra Kalisz, Anthony R. Bradley, and Charlotte M. Deane. Testing the limits of SMILES-based de novo molecular generation with curriculum and deep reinforcement learning. Nat Mach Intell, 5(4):386–394, April 2023. Number: 4 Publisher: Nature Publishing Group.

8. Ian J. Goodfellow, Jean Pouget-Abadie, Mehdi Mirza, Bing Xu, David Warde-Farley, Sherjil Ozair, Aaron Courville, and Yoshua Bengio. Generative Adversarial Networks, June 2014.

9. Xavier Bresson and Thomas Laurent. A Two-Step Graph Convolutional Decoder for Molecule Generation, June 2019. arXiv:1906.03412 [cs, stat] version: 2.

10. Nicola De Cao and Thomas Kipf. MolGAN: An implicit generative model for small molecular graphs, May 2018.

11. Chengxi Zang and Fei Wang. MoFlow: An Invertible Flow Model for Generating Molecular Graphs. In Proceedings of the 26th ACM SIGKDD International Conference on Knowledge Discovery & Data Mining, KDD ‘20, pages 617–626, New York, NY, USA, August 2020. Association for Computing Machinery.

12. Wengong Jin, Regina Barzilay, and Tommi Jaakkola. Hierarchical Generation of Molecular Graphs using Structural Motifs, April 2020. arXiv:2002.03230 [cs, stat].

13. Rebecca Boiarsky, Nalini Singh, Alejandro Buendia, Gad Getz, and David Sontag. A Deep Dive into Single-Cell RNA Sequencing Foundation Models. bioRxiv, page 2023.10.19.563100, October 2023. Publisher: Cold Spring Harbor Laboratory Section: New Results.

14. Fan Yang, Wenchuan Wang, Fang Wang, Yuan Fang, Duyu Tang, Junzhou Huang, Hui Lu, and Jianhua Yao. scBERT as a large-scale pretrained deep language model for cell type annotation of single-cell RNA-seq data. Nat Mach Intell, 4(10):852–866, October 2022. Number: 10 Publisher: Nature Publishing Group.

15. Christina V. Theodoris, Ling Xiao, Anant Chopra, Mark D. Chaffin, Zeina R. Al Sayed, Matthew C. Hill, Helene Mantineo, Elizabeth M. Brydon, Zexian Zeng, X. Shirley Liu, and Patrick T. Ellinor. Transfer learning enables predictions in network biology. Nature, 618(7965):616–624, June 2023. Number: 7965 Publisher: Nature Publishing Group.

16. Jiawei Chen, Hao Xu, Wanyu Tao, Zhaoxiong Chen, Yuxuan Zhao, and Jing-Dong J. Han. Transformer for one stop interpretable cell type annotation. Nat Commun, 14(1):1–14, January 2023. Number: 1 Publisher: Nature Publishing Group.

17. Haotian Cui, Chloe Wang, Hassaan Maan, Kuan Pang, Fengning Luo, and Bo Wang. scGPT: Towards Building a Foundation Model for Single-Cell Multi-omics Using Generative AI, July 2023. Pages: 2023.04.30.538439 Section: New Results.

18. Minsheng Hao, Jing Gong, Xin Zeng, Chiming Liu, Yucheng Guo, Xingyi Cheng, Taifeng Wang, Jianzhu Ma, L. Song, and Xuegong Zhang. Large Scale Foundation Model on Single-cell Transcriptomics, June 2023. Pages: 2023.05.29.542705 Section: New Results.

19. Graham Heimberg, Tony Kuo, Daryle DePianto, Tobias Heigl, Nathaniel Diamant, Omar Salem, Gabriele Scalia, Tommaso Biancalani, Jason Rock, Shannon Turley, Héctor Corrada Bravo, Josh Kaminker, Jason A. Vander Heiden, and Aviv Regev. Scalable querying of human cell atlases via a foundational model reveals commonalities across fibrosis-associated macrophages, July 2023. Pages: 2023.07.18.549537 Section: New Results.

20. Xiaodong Yang, Guole Liu, Guihai Feng, Dechao Bu, Pengfei Wang, Jie Jiang, Shubai Chen, Qinmeng Yang, Yiyang Zhang, Zhenpeng Man, Zhongming Liang, Zichen Wang, Yaning Li, Zheng Li, Yana Liu, Yao Tian, Ao Li, Jingxi Dong, Zhilong Hu, Chen Fang, Hefan Miao, Lina Cui, Zixu Deng, Haiping Jiang, Wentao Cui, Jiahao Zhang, Zhaohui Yang, Handong Li, Xingjian He, Liqun Zhong, Jiaheng Zhou, Zijian Wang, Qingqing Long, Ping Xu, The X.-Compass Consortium, Hongmei Wang, Zhen Meng, Xuezhi Wang, Yangang Wang, Yong Wang, Shihua Zhang, Jingtao Guo, Yi Zhao, Yuanchun Zhou, Fei Li, Jing Liu, Yiqiang Chen, Ge Yang, and Xin Li. GeneCompass: Deciphering Universal Gene Regulatory Mechanisms with Knowledge-Informed Cross-Species Foundation Model, September 2023. Pages: 2023.09.26.559542 Section: New Results.

21. Aditya Ramesh, Mikhail Pavlov, Gabriel Goh, Scott Gray, Chelsea Voss, Alec Radford, Mark Chen, and Ilya Sutskever. Zero-Shot Text-to-Image Generation, February 2021. arXiv:2102.12092 [cs].

22. Aditya Ramesh, Prafulla Dhariwal, Alex Nichol, Casey Chu, and Mark Chen. Hierarchical Text-Conditional Image Generation with CLIP Latents, April 2022. arXiv:2204.06125 [cs].

23. Carl Edwards, ChengXiang Zhai, and Heng Ji. Text2Mol: Cross-Modal Molecule Retrieval with Natural Language Queries. In Proceedings of the 2021 Conference on Empirical Methods in Natural Language Processing, pages 595–607, Online and Punta Cana, Dominican Republic, 2021. Association for Computational Linguistics.

24. Shengchao Liu, Yutao Zhu, Jiarui Lu, Zhao Xu, Weili Nie, Anthony Gitter, Chaowei Xiao, Jian Tang, Hongyu Guo, and Anima Anandkumar. A Text-guided Protein Design Framework, February 2023. arXiv:2302.04611 [cs, q-bio, stat].

25. Zhi Huang, Federico Bianchi, Mert Yuksekgonul, Thomas J. Montine, and James Zou. A visual–language foundation model for pathology image analysis using medical Twitter. Nat Med, 29(9):2307–2316, September 2023. Number: 9 Publisher: Nature Publishing Group.

26. Anna Gaulton, Louisa J. Bellis, A. Patricia Bento, Jon Chambers, Mark Davies, Anne Hersey, Yvonne Light, Shaun McGlinchey, David Michalovich, Bissan Al-Lazikani, and John P. Overington. ChEMBL: a large-scale bioactivity database for drug discovery. Nucleic Acids Res, 40(D1):D1100–D1107, January 2012. Publisher: Oxford Academic.

27. Aravind Subramanian, Rajiv Narayan, Steven M. Corsello, David D. Peck, Ted E. Natoli, Xiaodong Lu, Joshua Gould, John F. Davis, Andrew A. Tubelli, Jacob K. Asiedu, David L. Lahr, Jodi E. Hirschman, Zihan Liu, Melanie Donahue, Bina Julian, Mariya Khan, David Wadden, Ian C. Smith, Daniel Lam, Arthur Liberzon, Courtney Toder, Mukta Bagul, Marek Orzechowski, Oana M. Enache, Federica Piccioni, Sarah A. Johnson, Nicholas J. Lyons, Alice H. Berger, Alykhan F. Shamji, Angela N. Brooks, Anita Vrcic, Corey Flynn, Jacqueline Rosains, David Y. Takeda, Roger Hu, Desiree Davison, Justin Lamb, Kristin Ardlie, Larson Hogstrom, Peyton Greenside, Nathanael S. Gray, Paul A. Clemons, Serena Silver, Xiaoyun Wu, Wen-Ning Zhao, Willis Read-Button, Xiaohua Wu, Stephen J. Haggarty, Lucienne V. Ronco, Jesse S. Boehm, Stuart L. Schreiber, John G. Doench, Joshua A. Bittker, David E. Root, Bang Wong, and Todd R. Golub. A Next Generation Connectivity Map: L1000 Platform and the First 1,000,000 Profiles. Cell, 171(6):1437–1452.e17, November 2017. Publisher: Elsevier.

28. Jiangming Sun, Nina Jeliazkova, Vladimir Chupakhin, Jose-Felipe Golib-Dzib, Ola Engkvist, Lars Carlsson, Jörg Wegner, Hugo Ceulemans, Ivan Georgiev, Vedrin Jeliazkov, Nikolay Kochev, Thomas J. Ashby, and Hongming Chen. ExCAPE-DB: an integrated large scale dataset facilitating Big Data analysis in chemogenomics. Journal of Cheminformatics, 9(1):17, March 2017.

29. Hanjun Dai, Bo Dai, and Le Song. Discriminative Embeddings of Latent Variable Models for Structured Data, January 2020. arXiv:1603.05629 [cs].

30. Matthias Fey, Jan-Gin Yuen, and Frank Weichert. Hierarchical Inter-Message Passing for Learning on Molecular Graphs, June 2020. arXiv:2006.12179 [cs, stat].

31. Kristina Preuer, Philipp Renz, Thomas Unterthiner, Sepp Hochreiter, and Günter Klambauer. Fréchet ChemNet Distance: A Metric for Generative Models for Molecules in Drug Discovery. J. Chem. Inf. Model., 58(9):1736–1741, September 2018. Publisher: American Chemical Society.

32. Yusuf Roohani, Kexin Huang, and Jure Leskovec. Predicting transcriptional outcomes of novel multigene perturbations with GEARS. Nat Biotechnol, pages 1–9, August 2023. Publisher: Nature Publishing Group.

33. Prannay Khosla, Piotr Teterwak, Chen Wang, Aaron Sarna, Yonglong Tian, Phillip Isola, Aaron Maschinot, Ce Liu, and Dilip Krishnan. Supervised Contrastive Learning, March 2021. arXiv:2004.11362 [cs, stat].

34. Yonglong Tian, Dilip Krishnan, and Phillip Isola. Contrastive Multiview Coding. In Computer Vision – ECCV 2020: 16th European Conference, Glasgow, UK, August 23–28, 2020, Proceedings, Part XI, pages 776–794, Berlin, Heidelberg, August 2020. Springer-Verlag.

35. Aaron van den Oord, Yazhe Li, and Oriol Vinyals. Representation Learning with Contrastive Predictive Coding, July 2018.

36. Ming Ding, Zhuoyi Yang, Wenyi Hong, Wendi Zheng, Chang Zhou, D. Yin Junyang Lin, Xu Zou, Zhou Shao, Hongxia Yang, and Jie Tang. CogView: Mastering Text-to-Image Generation via Transformers, November 2021. arXiv:2105.13290 [cs].

37. Alec Radford, Jeffrey Wu, Rewon Child, David Luan, Dario Amodei, and Ilya Sutskever. Language Models are Unsupervised Multitask Learners.

38. Mike Lewis, Yinhan Liu, Naman Goyal, Marjan Ghazvininejad, Abdelrahman Mohamed, Omer Levy, Ves Stoyanov, and Luke Zettlemoyer. BART: Denoising Sequence-to-Sequence Pre-training for Natural Language Generation, Translation, and Comprehension, October 2019. arXiv:1910.13461 [cs, stat].

39. Colin Raffel, Noam Shazeer, Adam Roberts, Katherine Lee, Sharan Narang, Michael Matena, Yanqi Zhou, Wei Li, and Peter J. Liu. Exploring the Limits of Transfer Learning with a Unified Text-to-Text Transformer, July 2020. arXiv:1910.10683 [cs, stat].

40. OpenAI. GPT-4 Technical Report, March 2023. arXiv:2303.08774 [cs].

41. Rim Shayakhmetov, Maksim Kuznetsov, Alexander Zhebrak, Artur Kadurin, Sergey Nikolenko, Alexander Aliper, and Daniil Polykovskiy. Molecular Generation for Desired Transcriptome Changes With Adversarial Autoencoders. Frontiers in Pharmacology, 11, 2020.

42. TRIOMPHE: Transcriptome-Based Inference and Generation of Molecules with Desired Phenotypes by Machine Learning.

43. Jannis Born, Matteo Manica, Ali Oskooei, Joris Cadow, and María Rodríguez Martínez. PaccMannRL: Designing Anticancer Drugs From Transcriptomic Data via Reinforcement Learning. In Russell Schwartz, editor, Research in Computational Molecular Biology, Lecture Notes in Computer Science, pages 231–233, Cham, 2020. Springer International Publishing.

44. Tiago Pereira, Maryam Abbasi, Rita I. Oliveira, Romina A. Guedes, Jorge A. R. Salvador, and Joel P. Arrais. Deep generative model for therapeutic targets using transcriptomic disease-associated data—USP7 case study. Brief Bioinform, 23(4), July 2022. Publisher: Oxford Academic.

45. Matthias Fey and Jan Eric Lenssen. Fast Graph Representation Learning with PyTorch Geometric, March 2019.

46. Gregory Landrum. RDKit: Open-source cheminformatics. Release 2014.03.1, May 2014.

47. Abubakar Abid, Ali Abdalla, Ali Abid, Dawood Khan, Abdulrahman Alfozan, and James Zou. Gradio: Hassle-Free Sharing and Testing of ML Models in the Wild, June 2019.

48. Adam Paszke, Sam Gross, Francisco Massa, Adam Lerer, James Bradbury, Gregory Chanan, Trevor Killeen, Zeming Lin, Natalia Gimelshein, Luca Antiga, Alban Desmaison, Andreas Köpf, Edward Yang, Zach DeVito, Martin Raison, Alykhan Tejani, Sasank Chilamkurthy, Benoit Steiner, Lu Fang, Junjie Bai, and Soumith Chintala. PyTorch: An Imperative Style, High-Performance Deep Learning Library, December 2019.

